# Anticancer and antibacterial activities of secondary bioactive compounds from endosymbiotic bacteria of aphid and their surrounding organisms (Predator and protector)

**DOI:** 10.1101/2022.10.20.513013

**Authors:** Taghreed A. Alsufyani, Najwa Al-Otaibi, Noura J. Alotaibi, Nour H M’sakni, Eman M. Alghamdi

## Abstract

Secondary metabolites of bacterial origin are a valuable source of diverse molecules with antibacterial and anticancer activities. In the current study, 10 endosymbiotic bacteria were isolated from aphids, aphid predators and ants. Bacterial strains were identified based on the 16S rRNA gene. Crude extracts were prepared from each isolated bacteria and tested for their antibacterial activities using the disk diffusion method. The extracts of three bacterial species; *Planococcus* sp, *Klebsiella aerogenes, Enterococcus avium* from *Aphis punicae, Chrysoperia carnea* and *Tapinoma magnum*, respectively were found to have strong antibacterial activities against one or more of the five pathogenic bacteria tested. The inhibition zones ranged from 10.00± 0.13 to 20.00± 1.11mm with minimum inhibitory concentration (MIC) values ranged from 0.156 mg/mL to 1.25 mg/mL. The notable antibacterial activity was for the crude extract of *K. aerogenes* against *Klebsiella pneumonia* and *Escherichia coli* at MIC value of 0.156 mg/mL. The cytotoxic activity of the crude extracts varied according to the tested cell line. The most cytotoxic effect was for the extracts of *K. aerogenes* and *E. avium* at a concentration of 0.16 mg/mL against lung carcinoma epithelial cells (A549) with cell reduction of 79.4% and 67.2%, respectively. Crude extracts of *K. aerogenes* and *Pantoea agglomerans* at a concentration of 0.16 mg/mL showed 69.4% and 67.8% cell reduction against human colon cancer (Hct116), respectively. Gas chromatography–mass spectrometry (GC-MS) analysis of three crude extracts revealed the presence of several bioactive secondary metabolites that have been reported previously to possess antibacterial and anticancer properties. To the best of our knowledge, this is the first study to report the biological activities of endosymbiotic bacteria from aphids, aphid predators and ants. The promising data presented in this study opens a new avenue for alternative drugs to overcome the continuous emergence of multidrug-resistant bacteria and to find alternative drugs for traditional cancer therapies.

## Introduction

Cancer diseases and antibiotic-resistant phenomenon are two common challenges to the public health sectors. Different cancers are responsible for millions of annual deaths worldwide [1, 2]. Traditional therapeutic approaches such as surgery, radiotherapy, and chemotherapy are accompanied with harmful consequences such as anemia, alopecia, hormonal fluctuations, gastrointestinal mucositis, and brachial plexopathy [3]. Such deleterious effects pushed toward finding alternative treatment approaches [4]. Similarly, antibiotic-resistant bacteria continue to emerge despite discovering and producing new antibiotics [5]. Infections with Multidrug-resistant (MDR) bacteria usually result in serious disease complications, hospitalization, and deaths which pose challenges to public health authorities worldwide. Therefore, it is urgent to discover new antibiotics to cope with such antibiotic resistance phenomenon.

Endophytic and endosymbiotic bacteria represent an outstanding source of bioactive molecules with anticancer and antimicrobial activities [6]. Bacterial endophytes are less attention compared to fungal endophytes due to the lower yield concentration of crude extracts for the former [7, 8]. The review of Abdelghani et al, provides information about the crude extracts from bacteria that can be used as antibacterial and anticancer treatments [9]. It was estimated that *Actinomycetes* are responsible for the production of 70% of microbial secondary compounds while *Bacilli* and other bacteria produce around 7% and 1–2%, respectively [11]. Prodigiosin, methanolic pigment daptomycin, daptomycin and 3-benzyl-hexahydropyrrolo[1,2-a] pyrazine-1,4-dione are an example of secondary metabolites that are used as antibacterial compounds which extracted from *Serratia marcescens* [12], *Micrococcus* sp. [13], *Streptomyces roseospours* [14] and *Exiguobacterium indicum* [15], respectively. While anthracyclines, peptides, aureolic acids, and antimetabolites produced by *Actinomycetes* [16] and bleomycin from *Streptoalloteichus hindustanus* [17] are also reported to be cytotoxic against several cancer cells. Isolation of the secondary metabolites from natural sources (e.g., insects such as aphids) could be an alternative source of antimicrobial and anticancer chemotherapeutics that would help in reducing problematic infections affecting human health.

Aphids (Hemiptera: Aphididae) are a group of insects of agricultural importance which feed on many plant species. Most aphids establish mutualistic relationships with endosymbiotic bacteria, mainly known as obligates (i.e., primary) and facultative, that are housed in specialized cells called bacteriocytes [18, 19]. *Buchnera aphidicola* is a primary model of the obligate symbiont and can be found in almost all aphid species within three clades: *Aphidinae, Lachninae*, and *Fordini* [18, 20]. *B. aphidicola* plays an essential role in providing amino acids and nutrients lacking in the aphid diet [20]. Nine facultative symbionts, *Serratia symbiotica, Hamiltonella defensa, Regiella insecticola, Rickettsia, Rickettsiella, PAXS, Spiroplasma, Wolbachia, Arsenophonus*, have been shown positive effects on their hosts such as aphid fitness, immune pathway function and responses to natural enemies (e.g., defense against parasitoid attacks) [21, 22] and environmental stress (e.g., adaptation to thermal stress) [23-26]. However, much less attention has been given to the biological activities of these endosymbionts. Recent studies have shown that antibacterial and antifungal activity in the gall tissue of aphid [27, 28] or antimicrobial peptide against Pea aphid as bio-insecticides [29, 30]. To the best of our knowledge, no studies were conducted to investigate the antibacterial and anticancer activities of endosymbiotic bacteria of aphids, especially in Taif city, Saudi Arabia. Only a few attempts have studied the morphological identification of aphids in Taif [31] or controlling aphids infesting of Taif rose [32].

Here, we hypothesize that the crude extract from endosymbionts bacteria of aphids do present antimicrobial and anticancer activities. To test these hypotheses, we first isolated endosymbiotic bacteria associated with collected aphids from pomegranates, grapes and Taif rose along with aphid predators (ladybug and larvae of lacewing) and ants, and then we investigated the significant biological activities of the crude extracted against five strains of pathogenic bacteria (*S. aureus, S. epidermidis, E-coli, K. pneumoniae* and *E. cloacae)* and their anticancer activities against two cancer cell lines (adenocarcinoma human alveolar epithelial cells and human colon carcinoma) by Gas Chromatography-Mass Spectroscopy Analysis (GC-MS).

## 2. Materials and methods

### 2.1. Aphid sampling and identification

Aphids were collected from different plant hosts including pomegranate, grapes, Taif rose. Aphid predators such as ladybug, larvae of lacewing, and ants were collected from different sites located in Taif, Saudi Arabia. Sampling was collected from May to September of 2021 (Table 1). For molecular identification of the collected insects, DNA extraction was accomplished using the QIAamp® DNA Mini Kit (QIAGEN, Germany) as prescribed elsewhere [33]. The mitochondrial cytochrome oxidase gene (*COI*) was amplified by PCR using the primer set (LCO1490) (F-5’-GGTCAACAAATCATAAAGATATTGG-3’) and (R HCO2) (R-5’-TAAACTTCAGGGTGACCAAAAAATCA-3’) [34]. The PCR reaction was performed as described in our previous study [33]. PCR products were visualized on 1.5 % agarose gel using the BDA gel documentation system (Biometra-Germany). PCR products corresponding to the size of the amplified COI gene were retrieved from the gel and DNA sequencing was performed on both strands using a 3130xl Genetic Analyzer (Biosystems; Thermo Fisher Scientific, USA). The raw sequence data were edited and assembled using Bioedit software, version 7.2.5 (Ibis Biosciences, Carlsbad, Calif., USA) and the EditSeq program of the Lasergene software package, version 3.18 (DNAStar, Madison, Wisc., USA). The BLASTtool was used to identify the assembled sequences [35]. The assembled sequences were deposited in GenBank with the following accession numbers (aphids: MZ091377, MZ091379, and OL823183; aphid predators and ant: ON149796 to ON149799).

**Table 1:**
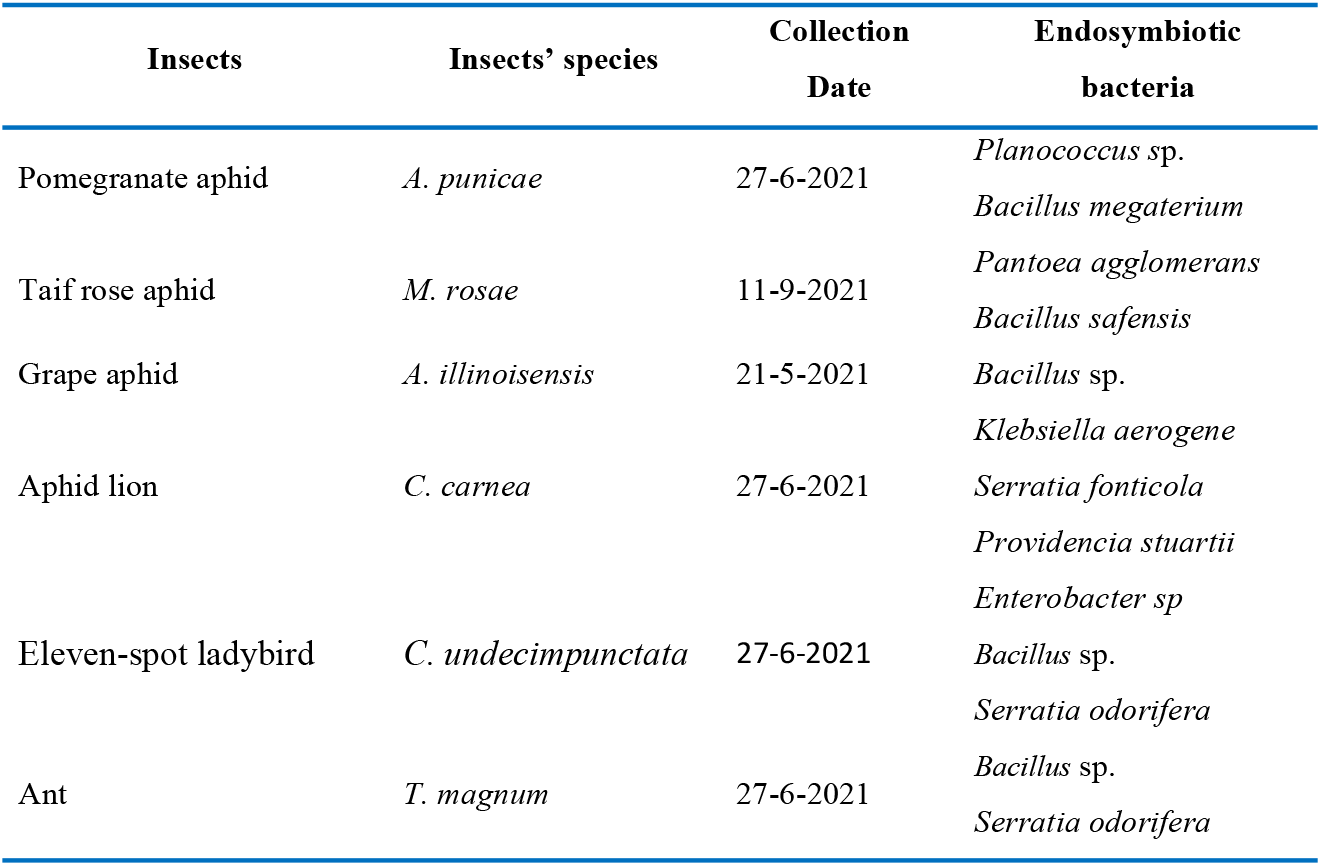
Insects samples, date collection and collection sites in Taif Governorate

### 2.2. Isolation of insect’s bacteria

Bacteria were isolated from identified three aphid species and their two species of predators and one species of ant. Insects were surface-sterilized with 70% ethanol for 3 min and washed three times with sterile PBS to get rid of surface contamination. Insects were then crushed and homogenized individually in nutrient broth media. Each isolation procedure was done in triplicate for each cultivar. Each triplicate suspensions were diluted individually (10^−1^ through 10^−5^). 100 μl of each dilution was plated on nutrient agar plates. Plates were incubated in an inverted position for 2–3 days at 30 °C. Growing colonies were picked up, then two rounds of purification were applied using nutrient agar plates. Purified bacterial isolates were picked, inoculated into 5 ml nutrient broth, and then incubated overnight at 37°C. After that, centrifugation was applied, the supernatant was removed, and the pellets was moved for the DNA extraction step.

### 2.3. DNA extraction, amplification and *16S rRNA* gene sequencing

DNA was extracted individually from isolated bacteria using DNeasy® Blood & Tissue kit (QIAGEN) kit, following the manufacturer’s instructions. V3 and V4 regions of the 16S rRNA gene was amplified using the following primer set Bakt_341F (CCTACGGGNGGCWGCAG), and Bakt_805R (GACTACHVGGGTATCTAATCC) [36]. The PCR reaction mixture contained 4 μLl of FIREPol® Ready to Load Master Mix (Solis BioDyne), 0.6 μl of each primer, 2 μl of isolated DNA, and water to bring the total volume to 20 μl. The PCR reaction conditions were initial denaturation at 95°C for 5 min, followed by 30 cycles of denaturation at 95°C for 40s, annealing at 55°C for 2 min and extension at 72°C for 1 min and a final extension step at 72°C for 7 min. The resultant PCR products were analyzed on 1.5% agarose gels, and visualized using the BDA gel documentation system (Biometra-Germany). PCR bands corresponding to the *16S rRNA* gene were excised from the gel, and purified by Illustra GFX PCR DNA and a gel band purification kit (GE Healthcare). DNA sequencing was accomplished using 3130xl Genetic Analyzer (Biosystems; Thermo Fisher Scientific, USA). Raw sequence data were edited and assembled using Bioedit software, version 7.2.5 (Ibis Biosciences, Carlsbad, Calif., USA), and the EditSeq program of the Lasergene software package, version 3.18 (DNAStar, Madison, Wisc., USA). Identification of bacterial isolates was achieved using the BLAST tool [35] and the *16S rRNA* gene sequences genes were deposited in the GenBank with the following accession numbers (OP320676 - OP320682).

### 2.4. Preparation of crude extract from endosymbiotic bacteria

Among the identified bacterial isolates, 10 isolates (Table 1) were selected to investigate their antibacterial and anticancer activities. Each bacterium was inoculated into Erlenmeyer flask containing 1L of nutrient broth and incubated in shaker incubator for 3-7 days at 200 rcf and 30°C. After seven days of incubation, bacterial cultures were centrifuged and sterile Amberlite® XAD16 (60 g/L; Sigma, BCBR6696V) was added to the supernatant and was shaken overnight at 200 rcf. The resin from each culture was collected individually after cheesecloth filtration into Erlenmeyer flasks, 300 ml of methanol was added to each flask and stirred for 2h. The methanol-soluble fraction was completely dried using a rotary evaporator, weighted, and redissolved in a suitable volume of methanol to unify the concentration of extracts.

### 2.5. Antibacterial activities of the crude extract against pathogenic bacteria

The effect of each endosymbiont crude extract was tested against five pathogenic bacteria, *Staphylococcus aureus* (ATCC6538), *Staphylococcus epidermidis* (ATCC14990), *Escherichia coli* (ATCC10536), *Klebsiella pneumoniae* (ATCC10031) and *Enterococcus cloacae* (ATCC13047), using the disk diffusion method [37]. The acetone-soluble crude extracts were filtered through a Millipore filter. Sterile filter paper discs with 8 mm in diameter were loaded with three different concentrations of each extract: 2, 5, and 10 mg. The discs were allowed to dry at room temperature and placed over Mueller-Hinton Agar plates seeded with the pathogenic bacteria. Disks loaded with ampicillin and tetracycline were used as positive control at a concentration of 30μg (Thermo Scientific, Applied Biosystems, Invitrogen, Gibco). The plates were incubated for 2h at -4°C to allow diffusion of the crude extracts and then transferred to the incubator at 37°C for 24h. The inhibition zones around the discs were measured and were considered as markers for antibacterial activity.

### 2.6. Resazurin-based 96-well plate microdilution assay for the determination of MICs

The principle of this assay is based on the reduction of resazurin by living bacteria resulting in colour changes from blue to pink and finally to colourless due to oxygen deficiency in the medium. A slight colour change indicates the inability of bacteria to grow and hence the antibacterial activity of the tested substance. Endosymbiont crude extracts that showed antibacterial activity was manipulated further to determine their minimum inhibitory concentration (MICs) and minimum bactericidal concentration (MBCs) using a microdilution assay. Serial dilutions from each extract were prepared in MH broth starting from 10 down to 0.039 mg/ml. The inoculum (100 μl) from each pathogenic bacterium were added to the assigned wells and the plate was incubated at 37°C for overnight. In the next day, 10 μl of resazurin sodium salt dye solution (0.02% w/v) was added to the assigned wells and the plate was incubated for 2 hr. After incubation, the plate was visually checked for colour change and wells with a known concentration showing slight colour change was determined as the MIC. The experiment was carried out in duplicate for each concentration of the crude extracts. The concentration above the MIC value was considered as that of MBC.

### 2.6. Antiproliferative activity of the endosymbiont crude extracts

The biological activity of the bacterial extracts against two types of cancer cell lines, adenocarcinoma human alveolar epithelial cells (A549), and human colon carcinoma (hct116), were tested in vitro using 3-(4,5-Dimethylthiazol-2-yl)-2,5-Diphenyltetrazolium Bromide (MTT) colorimetric assay. The cells were maintained in Dulbecco’s Modified Eagle’s Medium (DMEM) supplemented with 10% fetal bovine serum, 2 mM L-glutamine, and 1% Penicillin/Streptomycin. The test was performed in 96-well plates and the cells (5×10^4^ cells/ml) were seeded into the assigned wells and were incubated overnight at 37°C, 5% CO_2_, and 99% humidity. Two-fold serial dilutions of the crude extracts starting from 100 μg/ml, down to 3.125 μg/ml, were prepared in DMEM and added to the assigned wells. Control wells received methanol only and the plates were incubated for four days. The MTT was added to the cells, and the plates were incubated under the growth conditions for two hours. The medium was then removed and the cells were washed two times in phosphate buffered saline (PBS). The formed formazan crystals were dissolved in Isopropanol. The experiment was performed in duplicate for each concentration, the absorbance was measured at 595 nm and the average values obtained were considered. The following formula was used to calculate cell viability (Equation 1) [38]:

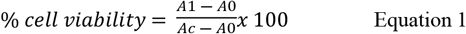

where A0, Ac and A1 are respectively the absorbance of blank, control solution and the extract at 595 nm. The IC_50_ % cell viability values were determined from % cell viability and the concentration curve according to the following equation 2 [39]:

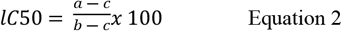

where a, b and c are respectively the absorbance at each concentration of the anticancer reagent, the absorbance at 0 μM of the anticancer reagent, and the absorbance of the blank.

### 2.7. Gas Chromatography–Mass Spectroscopy Analysis (GC-MS)

The chemical constituents of three crude extracts with significant biological activities against pathogenic bacteria and cancer cells were investigated using GC-MS (Shimadzu QP 2010 plus, Tokyo, Japan). The run was performed according to the protocol of [40]. The instrument has a silica column (5MS-RTX1) where pure helium gas was introduced, along with the methanolic extract of each sample, as a carrier at a fixed flow rate of 1 mL/min. The spectral detection was adjusted to the mass range of m/z 1.5 to 1000, and the electron ionization mode was applied. The scan sensitivity of the electron ionization was adjusted to a pictogram of octafluoronaphthalene (m/z 272, S/N > 200). Chemical compounds in each crude extract were identified based on their retention time (RT), percentage peak area (Equation 3), peak height, and molecular weight with the standard compounds of the databases of NIST and WILEY library to define the name and structure for each compound.

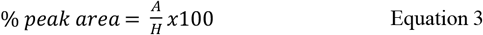

Where A, is the area of the target component (component A) peak and H, is the total area of all detected peaks to analyse quantity.

## 3. Results

### 3.1. Collection of insects and isolation of insects endosymbiotic bacteria

Three species of aphids were collected and identified molecularly based on sequence variation of the COI; *Aphis punicae, Macrosiphum rosae* and *Aphis illinoisensis* (Figure). The identified aphid predators used in this study included; *Chrysoperla carnea, Coccinella undecimpunctata*. In addition, one protective species; ants, *Tapinoma magnum*. Isolation of endosymbiotic bacteria from insects was attempted over nutrient agar plates. A total of 13 samples of bacteria were isolated and identified through the 16S rRNA gene sequences. The resultant sequences were introduced into the BLAST search tool of the gene bank resulted in several bacterial genera. As shown in Figure 1, phylogenetic analysis had confirmed the identification where related species are grouped into the same clade. Of the identified bacteria, only ten different species were selected to test their antibacterial and antiproliferative activities (Table 1).

**Figure 1.**
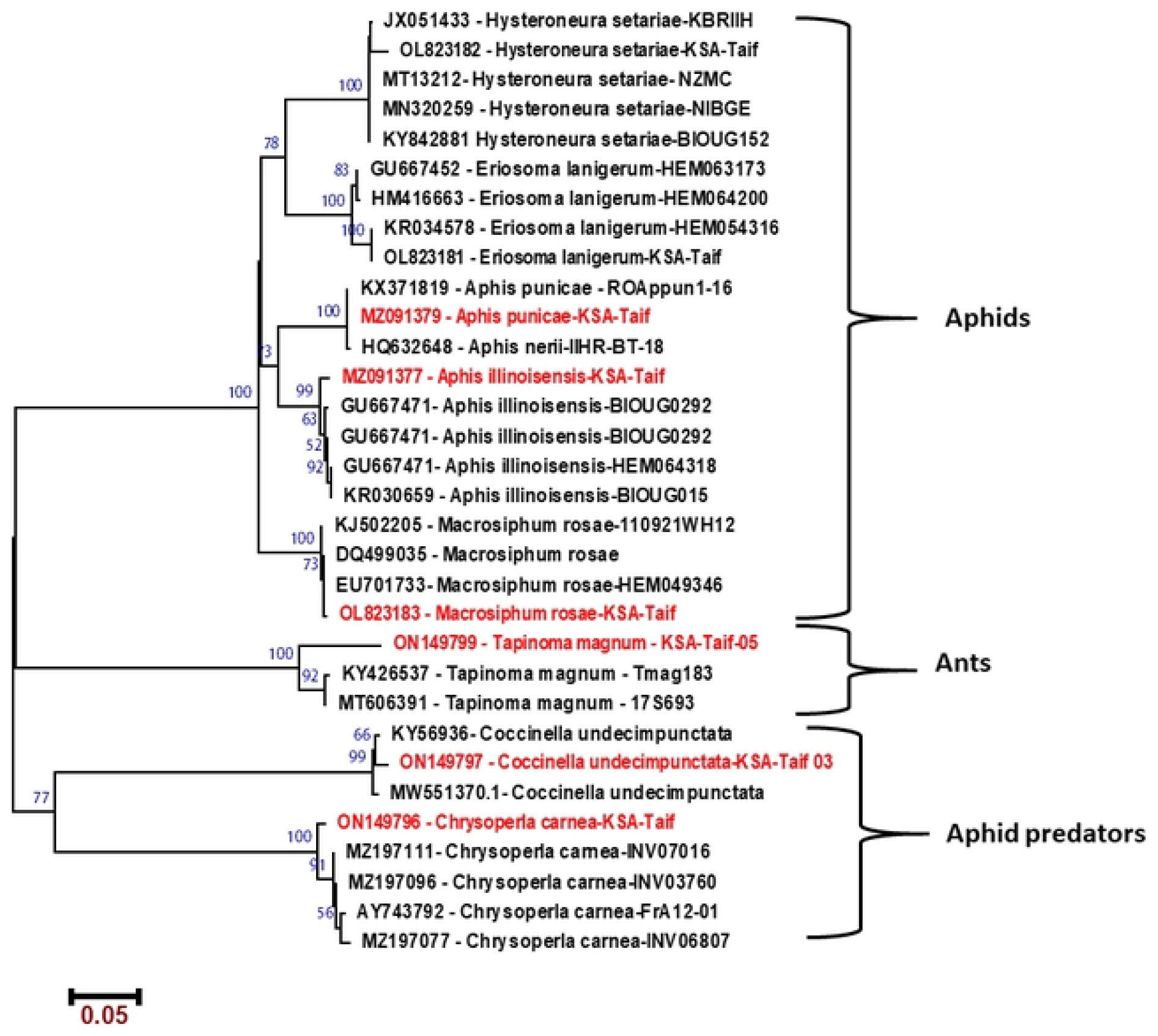
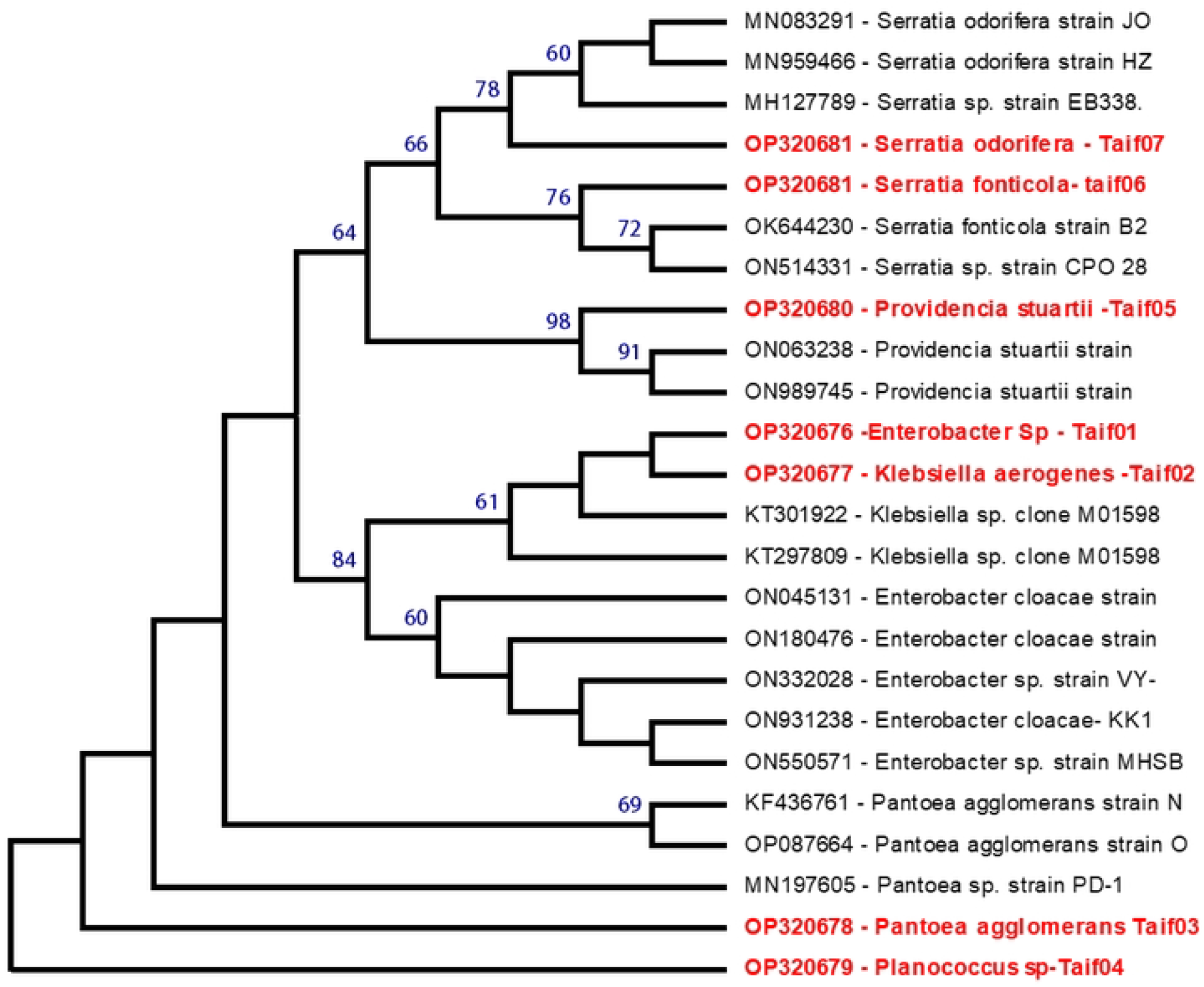
Neighbor-joining phylogenetic tree based on (A) mitochondrial COX1 gene of insects used in our study and other similar species from the Genbank database, and (B) 16S rRNA gene sequence of the endosymbiotic bacteria isolated from insects and other similar species retrieved from GenBank database. Insects from which bacteria were isolated and 16s rRNA identified bacteria are in red fonts. Numbers above the nodes refer to the bootstrap values generated after 1000 replications.

### 3.2. Antibacterial activities of endosymbiotic bacteria

No antibacterial activities were recorded for crude extracts after three days of incubation. However, after sept days of incubation, crude extracts of three endosymbionts; *Planococcus s*p, *Klebsiella aerogenes, Enterococcus avium* from *A. punicae, C. carnea* and *T. magnum* respectively were effective against both Gram-positive and Gram-negative bacteria with inhibition zones ranging from 10 to 20 mm. The effect of these crude extracts was comparable to the effect of tetracycline against some tested pathogenic bacteria. Extracts of *Enterobacter* sp, *Serratia odorifera, Pantoea agglomerans*, and *Bacillus megaterium* had no antimicrobial activities against the tested pathogenic bacteria. Crude extract of *Klebsiella aerogenes* was the most effective against four pathogenic bacteria; *S. aureus, S. epidermidis, K. pneumoniae*, and *E. coli* with inhibition zones of 15.00 ± 0.41, 16.00 ± 0.29, 20.00 ± 1.11, and (18.00 ± 0.65mm), respectively. Crude extract of *Planococcus* sp was effective against Gram-positive bacteria; *S. aureus* (17.00± 0.75mm) and *S. epidermidis* (16.00± 0.91mm) with weak antimicrobial activity against *K. pneumoniae* (11.00± 0.35mm).

The crude extract of the endosymbiont *Bacillus safensis* displayed a moderate antibacterial activity against *S. aureus*. According to the MIC values, the extracts of *Planococcus s*p displayed the lowest MIC value (0.3125 mg/mL) against *S. aureus* and *S. epidermidis*. Similarly, crude extract of *Enterococcus avium* was effective against *S. epidermidis*, and *K. pneumoniae* at MIC value of 0.3125 mg/mL and extract *Klebsiella aerogenes* was effective against *S. epidermidis* at the same MIC value (Table 2). Interestingly, the three crude extracts displayed low MIC values comparable with ampicillin and the second reference antibiotic, tetracycline (Table 2).

**Table 2.** Antibacterial activities of the methanol fraction of endosymbiont crude extracts at 10 mg/mL of concentration and MIC values.

| Crude extracts |  | Pathogenic bacteria |  |  |  |  |
| --- | --- | --- | --- | --- | --- | --- |
|  |  | *S. aureus* | *S. epidermidis* | *E. cloacae* | *K. pneumoniae* | *E. coli* |
| *Planococcus sp* | Inhibition zone | 17.00± 0.75 | 16.00± 0.91 | - | 11.00± 0.35 | - |
|  | MIC | 0.3125 | 0.3125 |  | 0.625 |  |
|  | MBC | 0.3125 | 0.3125 |  | 1.25 |  |
| *Enterobacter sp* |  | - | - | - | - | - |
| *Serratia odorifera* |  | - | - | - | - | - |
| *Klebsiella aerogenes* | Inhibition zone | 15.00± 0.41 | 16.00± 0.29 | - | 20.00± 1.11 | 18.00± 0.65 |
|  | MIC | 0.625 | 0.3125 |  | 0.156 | 0.156 |
|  | MBC | 0.625 | 0.3125 |  | 0.625 | 0.3125 |
| *Serratia fonticola* |  | - | - | - | - | - |
| *Providencia stuartii* | Inhibition zone | 09.00± 0.22 | - | - | - | - |
|  | MIC | 1.25 |  |  |  |  |
|  | MBC | 2.5 |  |  |  |  |
| *Enterococcus avium* | Inhibition zone | 10.00± 0.13 | 17.00± 0.37 | 11.00± 0.39 | - | - |
|  | MIC | 1.25 | 0.3125 | 0.625 |  |  |
|  | MBC | 2.5 | 0.625 | 1.25 |  |  |
| *Pantoea agglomerans* |  | - | - | - | - | - |
| *Bacillus megaterium* |  | - | - | - | - | - |
| *Bacillus safensis* | Inhibition zone | 10.00± 0.45 | - | - | - | - |
|  | MIC | 1.25 |  |  |  |  |
|  | MBC | 2.5 |  |  |  |  |
| Ampicillin |  | 19.00± 0.52 | 16.00± 0.95 | 15.00± 0.37 | - | 11.00± 0.33 |
|  |  | 0.3125 | 0.625 | 0.625 |  | 0.625 |
| Tetracycline |  | 35.00± 0.54 | 33.00± 0.36 | 25.00± 0.22 | 29.00± 0.63 | 30.00± 0.54 |
|  |  | 0.039 | 0.039 | 0.078 | 0.078 | 0.039 |

290 Data presented in the table are means of three replicates SD (n = 3).

### 3.3. Antiproliferative activities of endosymbiotic bacteria

The cytotoxic activities of 10 endosymbiont crude bacterial extracts were tested against A549 and Hct116 using MTT assay. In general, the intense of cytotoxicity was directly proportional to the concentration of the extracts. A high concentration of the crude extracts (4.3 mg/mL) was effective in reducing the cell growth of A549 and Hct116 cells. The crude extract of *Providencia stuartii* displayed the lowest cytotoxic activity against Hct116 (34.1% cell reduction) at a concentration of 4.3 mg/mL. Whereas, the same concentration resulted in 50.1% cell reduction against A549 cells. The extract of *Klebsiella aerogenes* displayed the highest cytotoxic activity against A549 cells (92.9% cell reduction) at a concentration of 4.3 mg/mL. This is followed by the extracts of *Pantoea agglomerans* and *Enterococcus avium* where cell reduction was 90.9 and 90.0%, respectively. In case of Hct116 cells, extract of *Klebsiella aerogenes* at a concentration of 4.3 mg/mL showed 89.5% cell reduction followed by extracts of *Pantoea agglomerans* (87.8% cell reduction) and *Enterococcus avium* (86.7% cell reduction) (Figure 2a).

**Figure 2.**
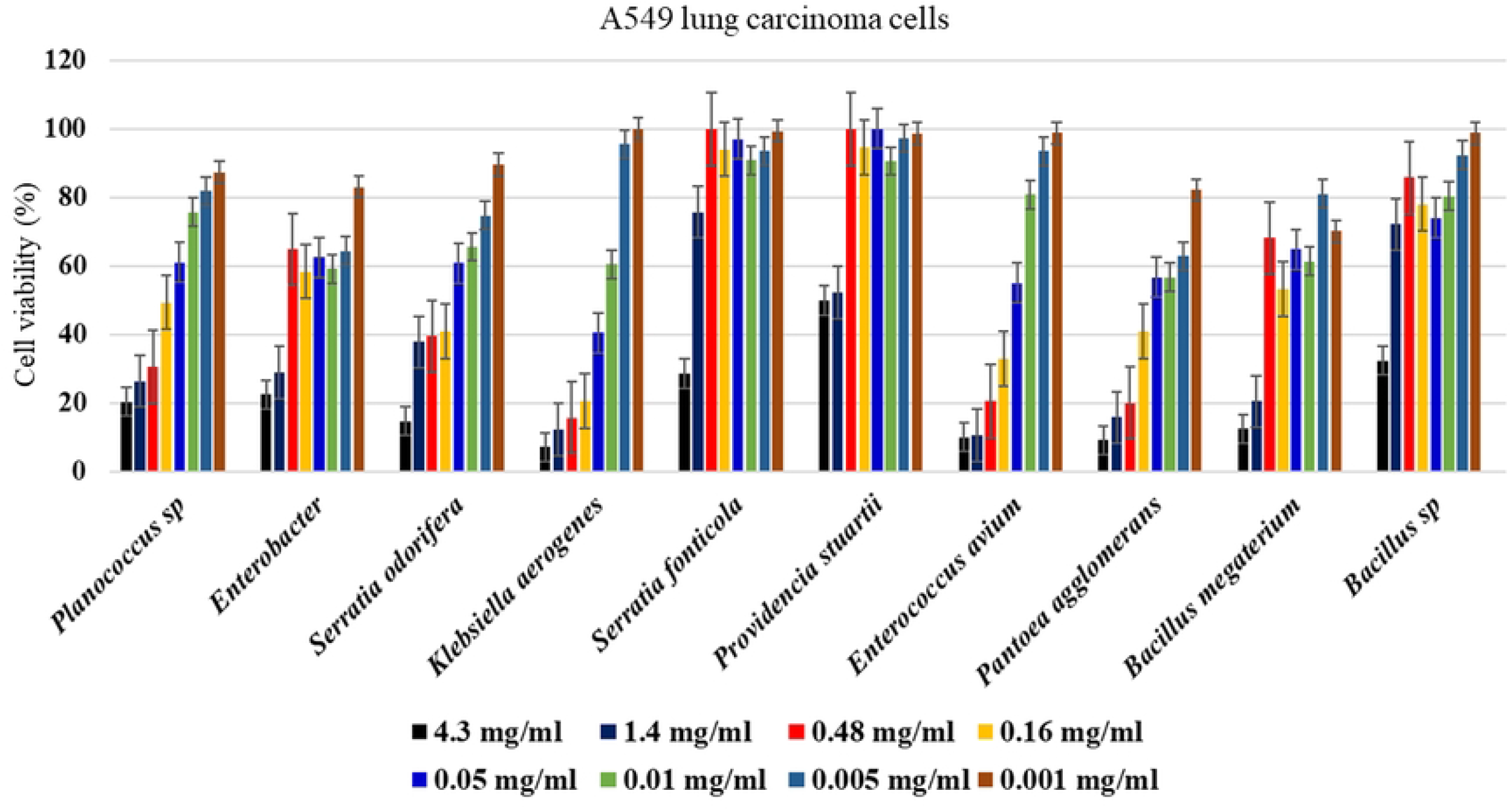
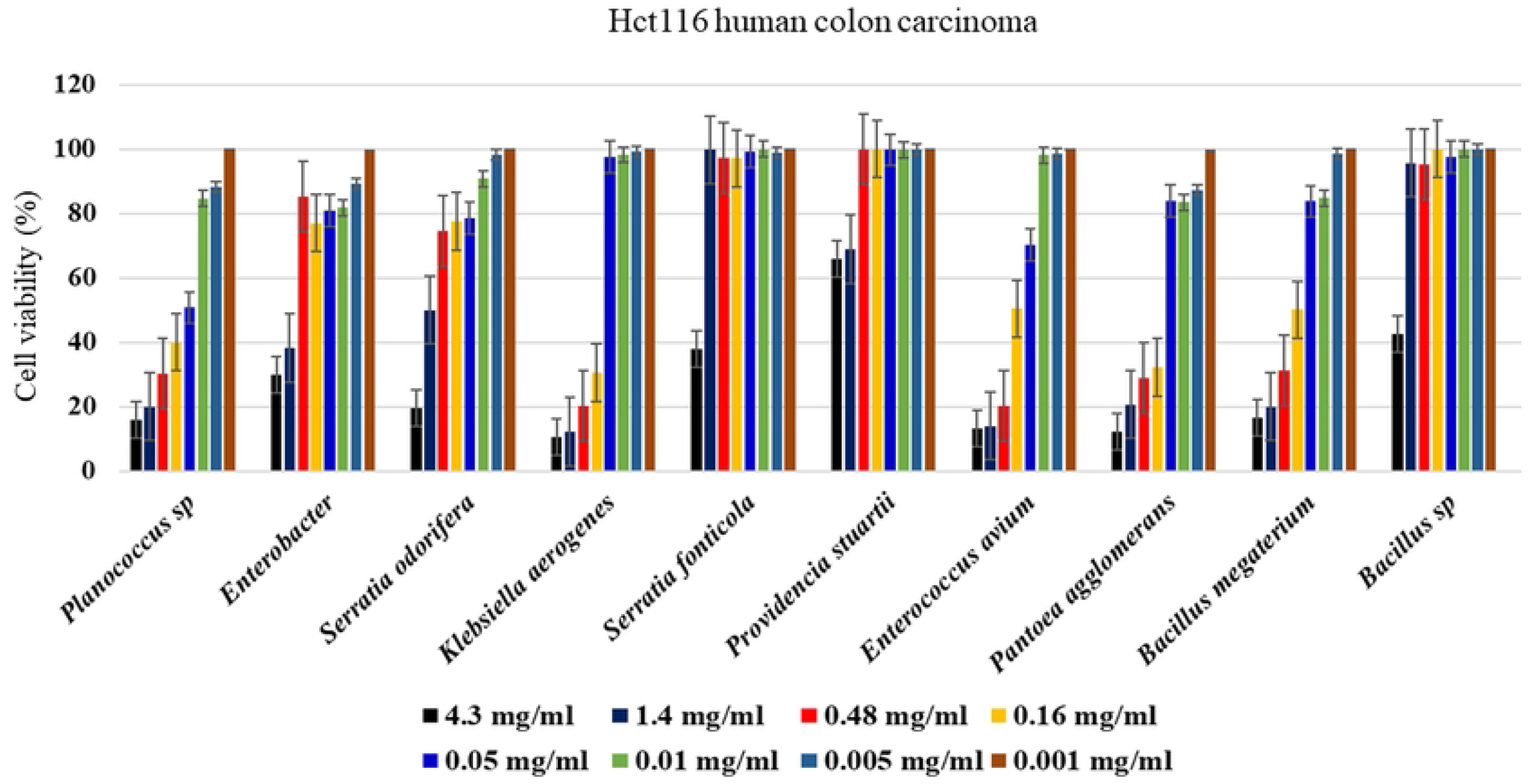
MTT cytotoxic activity assay of insect endosymbiotic bacterial crude extracts tested at different concentrations from 0.001 to 4.3 mg/mL, (A) A549 lung carcinoma cells and (B) Hct116 human colon carcinoma cells. Bars represent means ± standard deviations (SD) measured for each concentration (n=3)

At low concentration, 0.16 mg/mL, extracts of *Klebsiella aerogenes* and *Enterococcus avium* displayed the highest cytotoxicity against A549 cells with cell reduction of 79.4% and 67.2%, respectively. This is followed by the extracts of *Pantoea agglomerans* and *Serratia odorifera* where cell reduction was about 59.2%. Crude extracts of *Serratia fonticola* and *Providencia stuartii* had no impact on A549 cells above the concentration 1.4 mg/mL. For Hct116 cells (Figure 2b), extracts of *Klebsiella aerogenes* and *Pantoea agglomerans* at a concentration of 0.16 mg/mL showed 69.4% and 67.8% cell reduction, respectively. This is followed by extracts of *Planococcus sp* and *Bacillus megaterium* where cell reduction was 60% and 49.8% at the same concentration, respectively. At a concentration of 0.05 mg/mL, only the extract of *Planococcus sp* resulted in 49.8% cell reduction of Hct116 cells. In case of A549 cells (Figure 2a), extract of *Klebsiella aerogenes* and *Enterococcus avium* showed the highest cytotoxicity with cell reduction of 59.5% and 45%, respectively. The remaining extracts were ineffective or displayed a weak cytotoxicity against both cells at the same concentration. The LC_50_ values were determined for extract of *Planococcus sp* (0.16±8.22) mg/mL against A549 and (0.48±11.62) mg/mL against Hct116 cells. For the extract of *Klebsiella aerogenes*, LC_50_ values were (0.67±13.82) mg/mL and (0.38±9.20) mg/mL for A549 cells and Hct116 cells, respectively. The extract of *Enterococcus avium* showed the lowest LC_50_ value; (0.13±14.13) mg/mL and (0.22±11.13) mg/mL against A549 cell and Hct116 cells, respectively.

#### 2.9.2. Chemical analyses for endosymbiotic bacterial crude extracts

The GC-MS results have revealed several chemical compounds in each crude extract; 31 compounds in extract of *Planococcus sp*. 16 compounds in *Enterococcus avium* and 22 compounds in *Klebsiella aerogenes*. Among the major compounds identified in *Planococcus’s* insect origin crude extract are folinic acid, benzoic acid, 4-(1,1-dimethylethoxy),16-octadecadiynoic acid, methyl ester, Z-8-methyl-9-tetradecenoic acid, oleic acid, octadecanoic acid, 9,10-dihydroxy, and methyl ester. In case of *Enterococcus avium*, the major compounds are 13,16-octadecadiynoic acid, methyl ester, pyrrolo[1,2-a] pyrazine-1,4-dione, hexahydro-3-(2-methylpropyl), pentadecanoic acid, 8,11-octadecadiynoic acid, methyl ester, mono(2-ethylhexyl) phthalate; arachidonic acid; cholest-5-en-3-ol, and cholest-5-en-3-yl (9z)-9-octadecenoate. The crude extract of *Klebsiella aerogenes* has also a number of major compounds; pregnane-3,11,20,21-tetrol, cyclic 20,21-(butyl boronate), acetamide, n-(p-methoxybenzyl); hydrocinnamic acid, o-[(1,2,3,4-tetrahydro-2-naphthyl)methyl], 2,4-dimethylhexanedioic acid, dl-2,6-diaminoheptanedioic acid, oleic acid, didodecyl phthalate, and morphinan-4,5-epoxy-3,6-di-ol, 6-[7-nitrobenzofurazan-4-yl] amino. Some of the identified compounds were reported in the three crude extracts. The compounds structure, molecular weight and % peak area are shown in tables 3, 4 and 5.

**Table 3.** Chemical compositions for *Planococcus sp* crude extract

| Compound name | RT | %<br>Peak<br>area | Matching<br>factor | Molecular<br>formula | Molecular<br>weight |
| --- | --- | --- | --- | --- | --- |
| 2H-Pyran, 2-(2,5-hexadiynyloxy) tetrahydro | 4.10 | 13.72 | 683 | C<sub>11</sub>H<sub>14</sub>O<sub>2</sub> | 178 |
| dimethyl diphenyl tethylidyl pyrrolidine | 4.97 | 4.35 | 686 | C<sub>20</sub>H<sub>23</sub>N | 277 |
| 2-Aminoethanethiol hydrogen sulfate ester | 6.27 | 0.25 | 659 | C<sub>2</sub>H<sub>7</sub>NO<sub>3</sub>S<sub>2</sub> | 157 |
| 3-Methyl-4-nitro-5-(1-pyrazolyl) pyrazole | 6.48 | 0.36 | 651 | C<sub>7</sub>H<sub>7</sub>N<sub>5</sub>O<sub>2</sub> | 193 |
| 2(1H)-Naphthalenone, 4a,5,8,8a-Tetrahydro-4,4a-dimethyl | 8.00 | 0.38 | 628 | C<sub>12</sub>H<sub>16</sub>O | 176 |
| Folinic acid | 8.15 | 0.67 | 675 | C<sub>20</sub>H<sub>23</sub>N<sub>7</sub>O<sub>7</sub> | 473 |
| Benzoxazol, 2,3-dihydro-2-thioxo-3-diallylaminom ethyl | 9.14 | 0.14 | 695 | C<sub>14</sub>H<sub>16</sub>N<sub>2</sub>OS | 260 |
| 1-(á-d-Arabinofuranosyl)-4-O-difluor Omethyl uracil | 14.79 | 1.74 | 598 | C<sub>10</sub>H<sub>12</sub>F<sub>2</sub>N<sub>2</sub>O<sub>6</sub> | 294 |
| Benzoic acid, 4-(1,1-dimethylethoxy) | 15.05 | 0.47 | 686 | C<sub>11</sub>H<sub>14</sub>O<sub>3</sub> | 194 |
| 13,16-Octadecadiynoic acid, methyl ester | 19.09 | 1.48 | 733 | C<sub>19</sub>H<sub>30</sub>O<sub>2</sub> | 290 |
| 9-Octadecenoic acid, (2-phenyl-1,3-dioxolan-4-yl) methyl ester | 20.96 | 0.28 | 594 | C<sub>28</sub>H<sub>44</sub>O<sub>4</sub> | 444 |
| Leukotriene D4 methyl ester | 22.65 | 0.46 | 669 | C<sub>26</sub>H<sub>42</sub>N<sub>2</sub>O<sub>6</sub>S | 510 |
| Z-8-Methyl-9-tetradecenoic acid | 22.72 | 0.57 | 651 | C15H28O2 | 240 |
| Octadecanoic acid, 9,10-dihydroxy-, methyl ester | 23.38 | 0.34 | 610 | C19H38O4 | 330 |
| Dodecanoic acid, 2,3-bis(acetyloxy)propyl ester | 23.90 | 0.41 | 684 | C19H34O6 | 358 |
| Estra-1,3,5(10)-trien-17-ol | 24.80 | 0.89 | 672 | C18H24O | 256 |
| D-Fructose, diethyl mercaptal, pentaacetate | 25.17 | 0.55 | 622 | C20H32O10S2 | 496 |
| 7,8,15,16-Tetramethyl-1,9-dioxacyclohexadeca-4,13-diene-2,10-dione | 25.27 | 0.76 | 538 | C18H28O4 | 308 |
| D-fructose, diethyl mercaptal, pentaacetate | 25.58 | 0.67 | 627 | C20H32O10S2 | 496 |
| 6,9,12-Octadecatrienoic acid, phenylmethyl ester, (z,z,z) | 39.05 | 0.25 | 761 | C25H36O2 | 368 |
| Cholest-5-en-3-ol (3a) | 39.57 | 3.96 | 715 | C27H46O | 386 |

**Table 4.** Chemical compositions for *Enterococcus avium* crude extract

| Compound name | RT | % Peak area | Matching factor | Molecular formula | Molecular weight |
| --- | --- | --- | --- | --- | --- |
| Pregnane-3,11,20,21-tetrol, cyclic 20,21-(butyl boronate), (3à,5á,11á,20r) | 4.09 | 12.16 | 630 | C25H43BO4 | 418 |
| Cyclopropanedodecanoic acid, 2-octyl-, methyl ester | 14.77 | 1.15 | 636 | C24H46O2 | 366 |
| Methyl 16-hydroxy-hexadecanoate | 19.06 | 1.21 | 696 | C17H34O3 | 286 |
| 14-Methyl-, methyl esterhexadecanoic acid, methyl ester | 23.11 | 35.13 | 869 | C17H34O2 | 270 |
| N-(3-Chlorobenzylidene)-10-undecenoic acid hydrazide | 23.40 | 0.57 | 592 | C18H25ClN2O | 320 |
| Pyrrolo[1,2-a]pyrazine-1,4-dione, hexahydro-3-(2-methylpropyl) | 24.85 | 1.34 | 685 | C11H18N2O2 | 210 |
| Pentadecanoic acid | 24.95 | 0.94 | 629 | C15H30O2 | 242 |
| Oxacyclotetradecan-2-one, 13-methyl | 26.00 | 0.52 | 605 | C14H26O2 | 226 |
| 8,11-Octadecadiynoic acid, methyl Ester | 27.14 | 0.64 | 610 | C19H30O2 | 290 |
| 1,2-Benzenedicarboxylic acid | 33.57 | 0.79 | 782 | C24H38O4 | 390 |
| Mono(2-ethylhexyl) phthalate | 36.42 | 0.72 | 644 | C16H22O4 | 278 |
| Arachidonic acid | 39.41 | 0.32 | 661 | C20H32O2 | 304 |
| Cholest-5-en-3-yl (9z)-9-octadecenoate | 39.57 | 1.81 | 716 | C45H78O2 | 650 |

**Table 5.** Chemical compositions for *Klebsiella aerogenes* crude extract

| Compound name | RT | % Peak area | Matching factor | Molecular formula | Molecular weight |
| --- | --- | --- | --- | --- | --- |
| 2h-Pyran, 2-(2,5-hexadiynyloxy) tetrahydro | 4.09 | 9.07 | 662 | C11H14O2 | 178 |
| Acetamide, n-(p-methoxybenzyl) | 4.62 | 7.16 | 647 | C10H13NO2 | 179 |
| 4a,8a-(Methanothiomethano)nap hthalene, 1,4,5,8-tetrahydro- | 14.74 | 1.68 | 680 | C12H16S | 192 |
| 13,16-Octadecadiynoic acid, methyl ester | 19.07 | 1.31 | 769 | C19H30O2 | 290 |
| 2,4-Dimethylhexanedioic acid | 23.42 | 0.86 | 690 | C8H14O4 | 174 |
| Oleic acid | 24.87 | 3.05 | 722 | C18H34O2 | 282 |
| 9-Octadecenoic acid (z), methyl ester | 26.30 | 4.63 | 830 | C19H36O2 | 296 |
| Cyclopentaneundecanoic acid, methyl ester | 26.81 | 5.34 | 853 | C17H32O2 | 268 |
| Didodecyl phthalate | 33.58 | 0.68 | 847 | C32H54O4 | 502 |
| 2-Aminoethanethiol hydrogen sulfate (ester) | 39.57 | 2.14 | 721 | C2H7NO3S2 | 157 |

## Discussion

The continuous emergence of MDR bacteria and the deleterious effects of traditional cancer treatments directed researchers to seek for alternative approaches. In the past few decades, natural products secured a greater than 40% of the drugs approved as anti-microbial or anti-proliferative agents [41]. Bioactive compounds of natural origin have received much attention due to their safety profile, effectiveness, and availability. A huge body of research focuses mainly on medicinal plants and investigates their anticancer and antibacterial activities against a wide range of pathogenic bacteria and cancer cells [42-44]. Another important source of bioactive compounds is the endophytic bacteria [6, 45-49]. Few studies had investigated the antibacterial and anticancer activities of endosymbiotic bacteria. In these studies, endosymbiotic bacteria such as *bacilli* were isolated from many arthropods [50], *Enterobacter* sp from *Dysidea granulosa* [51] and *Bacillus brevis*, and *Bacillus choshinensis* from the earthworm *Pheretimasp* [52].

In the current study, crude extracts collected after three days of incubation displayed neither antibacterial nor anticancer activities. Strong biological activities against bacteria and cell lines was obtained from extracts collected after seven days of incubation. After seven days, the depletion of nutrients in the medium force bacteria to get into the stationary phase and release secondary metabolites. In several studies, secondary metabolites were obtained after seven days’ incubation period of bacteria [5, 48, 53]. Among the tested crude extracts, *K. aerogenes*’s extract displayed strong antibacterial activities against both Gram-positive and Gram-negative bacteria. Such antibacterial effect is greater than the crude extract of the *Raoultella ornithinolytica*, endophytic bacteria, against *E. coli* and *K. pneumoniae* where the MIC values were 0.5 and 0.25 mg/mL, respectively [5]. In another study, crude extract of earthworm endosymbiotic bacteria, *Bacillus* sp, resulted in an inhibition zone of 16.88 mm for *Staphylococcus aureus* [52].

In our study, crude extracts of *Planococcus* sp. and *K. aerogenes* inhibited the growth of *S. aureus* with inhibition zones of 17.0 and 15.0 mm, respectively. Crude extracts of *Bacillus amyloliquefaciens* isolated from a marine sponge showed strong antibacterial activity against *S. aureus* (inhibition zones 20.0 mm). In the same study, crude extract of *Alcaligenes faecalis* displayed high activity against *E. coli* (inhibition zone 16.0 mm) [54]. The extract of *Bacillus safensis* displayed a moderate activity against *S. aureus* (MIC 1.25 mg/mL) and its extract had no impact on the remaining tested bacteria (Table 2). In the study of Sebola et al, the extract of endophytic *B. safensis*s extract showed activity against *E. coli* at a concentration of 0.25 mg/mL [5].

In another study, *B. safensis* isolated from *Ophioglossum reticulatum* displayed antibacterial activity against *S. aureus* and *E. coli* with inhibition zones of 15.0 mm and 11.33 and 15.0 mm, respectively [55]. Such difference in antibacterial activity could be attributed to the host where bacteria, in some instances, share to some extent the chemistry of their hosts [56]. In the study of Sebola and Mukherjee, *B. safensis* was isolated from ethnomedicinal plants in which bacteria may mimic the chemistry of their host plants and produce more bioactive and effective chemicals [56]. The association of bacteria with aphids was reported to play several functions such as providing essential amino acids, reproductive manipulations, thermal adaptation [57]. In addition, chemicals produced by these symbiotic bacteria also help in defence against pathogenic microbes and predators [58-61].

For the antiproliferative activities, the data of MTT assay showed strong cytotoxicity of some crude extracts including those of *K. aerogenes* (LC_50_: 0.67±13.82 mg mL^-1^ for A549 cells and 0.38±9.2 mg/mL for Hct116 cells) *Pantoea agglomerans, Enterococcus avium* (LC_50_: 0.13±14.13 mg/mL for A549 and 0.22±11.13 mg/mL for Hct116 cells) and *Planococcus sp* (0.16±8.22 mg/mL for A549 cells and 0.48±11.62 mg/mL for Hct116 cells). Crude extract of endophytic *B. safenesis* displayed notable cytotoxicity against A549 cells with 50% cell reduction at a concentration of 100 *μ*g/mL [5]. In another study, crude extract of *B. safensis* isolated from sponges was reported to have cytotoxic activity against HepG2, HCT, and MCF7 [62]. However, in our study, crude extract of *B. safensis* showed weak or no cytotoxic activity against A549 and Hct116 cells at low concentrations (Figure 1a,b). Only at high concentration 4.3 mg/mL, the crude extract resulted in 67.7 and 57.8% cell reduction in A549 cells and Hct116, respectively. The difference in crude extract activity across the studies could be attributed the host from which the bacteria was isolated [56]. Interestingly, crude extract of *Bacillus* sp was able to reduce cell viability of A549 cells to 0% at 0.1 mg/mL [63]. The activity of *Bacillus* sp. against various types of cancer cells were reported [64-68]. These findings highlight the importance of *Bacillus* sp. As a source of biological active compounds against cancer cells.

The chemical constituents of three crude extracts of *Planococcus* sp, *Enterococcus avium*, and *K. aerogenes* were analysed using GC-MS. Many of the identified compounds (Tables 3-5) were reported in several studies to have biological activities against pathogenic bacteria and cancer cells. For example, 2H-pyran, 2-(2,5-hexadiynyloxy) tetrahydro, identified in the three extracts, was isolated previously from *Aspergillus terreus* and reported to have antibacterial and antifungal activity [69]. It was also identified in *P. guajava, Pogostemon quadrifolius*, and many other medicinal plants with different biological activities [70-72]. In a recent study, the pyran compound was reported to act against *Salmonella enterica* serovar Typhi [73]. The compound dimethyl diphenyl tethylidyl pyrrolidine from *Planococcus sp* was effective against several pathogenic bacteria such as *Porphyromonas gingivalis, S. aureus, S. pyrogenes* and *E. coli* and antifungal activity against *Aspergillus niger, Candida albicans* and *Aspergillus clavatus* [74, 75]. 3-Methyl-4-nitro-5-(1-pyrazolyl) pyrazole which was identified in the crude extract of *Planococcus* sp was reported in the literature to have antimicrobial activities against *B. subtilis, S. aureus, P. fluorescens, P. aeruginosa* and *E. coli*. In addition, it had anticancer activities against various types of cell lines [76-78].

Long chain fatty acids such as oleic acid, cholest-5-en-3-ol and were also identified in one or more of the 3 extracts. Oleic acid was reported to have antiproliferative activity against hepatocellular carcinoma cell lines, tongue squamous cell carcinomas and other cancer cells [79, 80]. In addition, it had antibacterial activity against *S. aureus* and *E. coli* [81]. Cholest-5-en-3-ol was also reported to have antitumor and antibacterial activities against several pathogens including *<u>Acinetobacter baumannii</u>* [82]. Other compounds that were identified and reported for their biological activities include mono (2-ethylhexyl) phthalate [83], pentadecanoic acid [84], acetamide n-(p-methoxybenzyl [85, 86].

In conclusion, the current study highlights the importance of endsymbiotic bacteria of aphids, aphid predators and ants as sources for bioactive compounds against cancer and pathogenic bacteria. Of the tested bacteria, crud extracts of *Planococcus* sp, *Klebsiella aerogenes, Enterococcus avium* were the most effective against pathogenic bacteria and cancer cell lines. Large-scale studies to investigate more endosymbiotic bacteria of aphids and other insects could potentially lead to the discovery of new bacterial species with potent anticancer and antimicrobial activities.

## Institutional Review Board Statement

Not applicable.

## Informed Consent Statement

Not applicable.

## Data Availability Statement

All data generated or analyzed during this study are included in this published article.

## Acknowledgments

The authors extend their appreciation to the Deputyship for Research & Innovation, Ministry of Education in Saudi Arabia, for funding this research work through the project number (1/441/119).

